## Supplementary figures for "KIR gene content imputation from single-nucleotide polymorphisms in the Finnish population"

**a** **b**

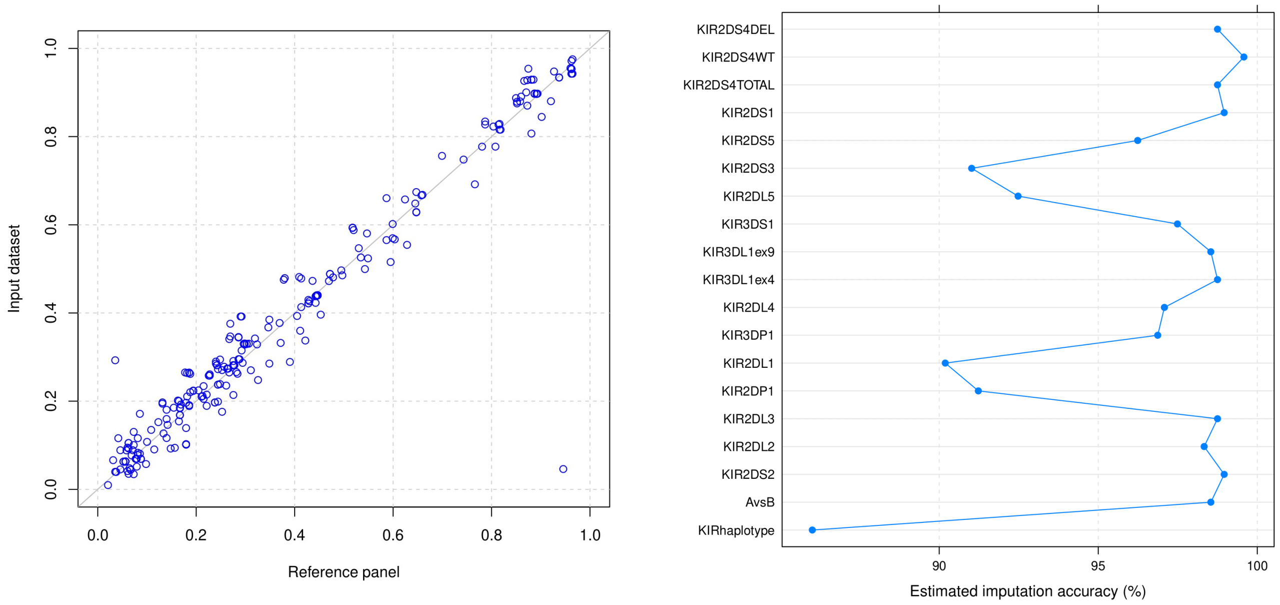


**Figure S1**. KIR*IMP input data validity check. **a)** Scatterplot showing the SNP allele frequencies in the KIR*IMP reference panel (x-axis) and in the input data (y-axis). **b)** KIR*IMP imputation accuracy estimate based on SNPs present in the input data.
